# Systematic identification of genomic non-response biomarkers to cancer therapies

**DOI:** 10.1101/2025.10.06.680777

**Authors:** Joseph Usset, Joep de Ligt, Sophie Roerink, Paul Roepman, Edwin Cuppen, Francisco Martínez-Jiménez

## Abstract

The costs of cancer therapies are rising rapidly across the globe, with novel therapies like targeted treatment and immunotherapies as major contributors, but their effectiveness can be low or uncertain due to limited post market surveillance. Reliable biomarkers to identify patients highly unlikely to respond to cancer therapies represent an increasingly important clinical and societal need, as they could prevent unnecessary treatments, reduce side effects, and alleviate pressure on healthcare systems. Here, we developed a robust statistical framework and applied it to whole-genome and transcriptome sequencing data of cancer patients (n = 2,596) with advanced disease. Our approach systematically identified known and potentially novel genomic and transcriptomic biomarkers of non-response, such as immune evasion driver events in skin melanoma patients treated with anti-PD-1 checkpoint inhibitors, and *KRAS*^G12^ mutations in metastatic colorectal cancer patients treated with different chemotherapy regimens. Despite the identification of these promising non-response signals, an analytical power analysis revealed that for most treatments and/or cancer types the cohort sizes are underpowered. Our results underscore the promises and the urgent need for expanding response-annotated real-world comprehensive genomics datasets to enable robust biomarker identification and validation.

## Main

Precision oncology has largely focused on the identification of targetable biomolecules that guide novel treatments, which primarily has revealed biomarkers for targeted therapies and immune checkpoint blockade (1,2). To our knowledge, however, relatively limited attention has been given to the identification of genomic biomarkers that identify patients unlikely to respond, so-called non-responder biomarkers, despite their substantial clinical value. The key distinction in evaluating non-response biomarkers is the robust evidence required to justify withholding an otherwise beneficial therapy. Clinically, this demands a high degree of confidence that the likelihood of benefit is negligible. Specifically, while a biomarker associated with a 30–40% response rate may be considered useful for identifying likely responders, a non-response biomarker must be linked to a response rate acceptably low, or ideally 0%, to be considered actionable in guiding treatment exclusion. When expected patient benefits are high (survival increased by years instead of weeks) or side-effects of the treatment are minimal, the bar for not starting a specific treatment will be higher than for treatments with limited effects and/or high risks of severe side-effects. By flagging individuals unlikely to benefit from specific treatments, non-responder biomarkers can spare patients unnecessary toxicity, reduce pressure on healthcare system (3,4), and prompt timely consideration of alternative strategies, including clinical trial enrolment.

Analytical studies are increasingly uncovering genomic determinants of treatment resistance across cancer types. For example, alterations in the WNT/β-catenin pathway and cyclin-dependent kinases (*CDK4/6*) have been implicated in limiting the efficacy of *EGFR* tyrosine kinase inhibitors (5), while elevated chromosomal instability scores have been associated with resistance to multiple standard chemotherapeutic agents (6). Similarly, a higher cumulative burden of oncogenic driver events or pathway-level alterations has been proposed as a marker of diminished response to later-line therapies (7). While such biomarkers are valuable for identifying patients at risk of treatment failure, they rarely quantify the probability of non-response, i.e. the consequences of withholding a therapy based on the identified biomarker. A notable example comes from the field of check-point inhibitors for non-small cell lung cancer (NSCLC) where a low tumor mutational burden (TMB) is associated with poor response. However, when the patient population is stratified based on a TMB cut-off of 10 mutations per Mb, the group of patients that would not receive treatment would still include 30% of responding patients (8). The work of Van de Haar et al. (9) shows that an acceptable stratification can be achieved when the TMB biomarker is combined with *STK11, KEAP1*, or *EGFR* alterations, resulting in a subgroup of 20% of all patients for which clinical benefit from immune checkpoint blockade was below 3%. While illustrative, this finding remains anecdotal and underscores a broader gap: the lack of systematic approaches to identify, validate, and evaluate the clinical implantation of non-response biomarkers across tumor types and therapeutic contexts.

The rapid implementation of comprehensive genomic profiling approaches for cancer diagnosis across institutions (10–12) coupled with harmonized clinical-genomic guidelines for reporting (13), present a unique opportunity to systematically identify non-responder biomarkers across diverse patient population.

In this study, we present an analytical framework to systematically identify genomic and transcriptomic biomarkers of non-response from real world clinical and molecular data. We analyzed data from 2,596 patients with annotated clinical outcomes in the Hartwig Medical Foundation cohort (of 7,010 total), across cancer types and treatment regimens. Our analysis uncovered multiple known and novel biomarkers associated with non-response to immune checkpoint blockade (ICB) and other cancer therapies that displayed strong signals in our discovery cohort. However, power analysis revealed for most treatment-tumor type combinations the existing cohorts remain underpowered to confidently identify biomarkers associated with extremely low response rates. These findings underscore the necessity for vastly increasing real world clinicogenomics datasets, for example by federated data-sharing approaches, to enhance detection capability and translate data-driven insights into patient-centered clinical decisions.

## Results

### A framework for non-responder biomarker identification

We developed an analytical framework to systematically identify biomarkers associated with non-response to cancer therapies using whole-genome (WGS) and whole-transcriptome (WTS) data. Non-responders were defined as patients who did not achieve at least a partial response or who had stable disease lasting less than six months based on RECIST v1.1 (14). We conducted a comprehensive analysis of clinicogenomic 1,157 features derived from the WiGiTS pipeline (15) to assess their association with non-response across a range of systemic and targeted cancer therapies, which were grouped by mode of action and/or target. Genomic features encompassed a diverse range of somatic features such as tumor mutation burden, copy number alterations, driver events, mutational signatures, genomic instability patterns, and immunogenomic factors, among others. Moreover, we incorporated 1,560 transcriptomic features derived from curated gene sets. **Supplementary Table 1** provides a detailed overview of all the biomarkers analyzed.

To identify biomarkers of non-response, we used the Hartwig Medical Foundation WGS/WTS cohort (n = 7,010), focusing on the subset of patients with post-biopsy treatment and clinical response data assessed via RECIST v1.1 (n = 2,596). We stratified patient groups according to primary tumor location, immunohistochemistry tumor molecular markers (e.g., ER+/HER2− breast cancer), and post-biopsy treatments, based on different levels of treatment annotation granularity (drug name, mechanism of action and treatment type). In total, we defined 56 treatment cohorts, each comprising at least 30 patients with a minimum of 15 non-responders and responders (**Supplementary Figure 1, Supplementary Table 2**). Most of the cohorts had limited size with median counts equal to 62 patients, and median number of non-responders equal to 36 patients.

### Systematic identification of non-response biomarkers

To systematically identify features associated with non-response, we screened for biomarkers significantly enriched in patient subgroups with low response rates (<5 %) and assessed statistical significance by performing both Fisher’s exact tests and univariate Cox proportional hazards analyses across all features within each defined cohort (**Supplementary Figure 1a**).

Applying a stringent multiple-testing corrected non-response screen—defined as an observed objective response rate below 5% and FDR-adjusted *P* < 0.10 —we identified only two biomarkers that were significantly associated with non-response across treatment cohorts using both models **(Figure 1, Supplementary Figure 2)**. First, low to moderate RNA-expressed neoantigen load was associated with non-response in NSCLC patients treated with anti–PD-1 therapy (4% response rate; 23 non-responders; 1 responder; baseline response rate 49%; odds ratio = .048; Fisher adjusted p-value = .052, Cox-PH adjusted p-value = .087). Second, the group of *KRAS*^*G12*^ mutations—previously implicated in resistance to trifluridine/tipiracil in metastatic colorectal cancer (mCRC) (16) —was associated with non-response across multiple chemotherapy regimens in colorectal cancer (5% response rate; 3 responders out of 65 patients; baseline response rate 23%; odds ratio = .072; Fisher adjusted p-value = .093, Cox-PH adjusted p-value = .011). Moreover, within the *KRAS*^*G12*^ mutations, the canonical *KRAS*^*G12D*^ hotspot showed 23 non-responders and 0 responders within the mCRC cohort (Fisher adjusted p-value = .36, Cox-PH adjusted p-value = .015). Two additional non-response biomarkers reached statistical significance only in Fisher’s exact test. These included immune evasion driver events—primarily biallelic *B2M* loss—in melanoma patients treated with immune checkpoint blockade (0% response rate; 0 responders out of 11 patients; baseline response rate 51%; odds ratio = 0; Fisher adjusted p-value = .098, Cox-PH adjusted p-value = .16) (17,18)(**Supplementary Note** provides further validation using cBioPortal (19–22)), and elevated complement and coagulation cascade RNA expression in prostate cancer patients undergoing androgen deprivation therapy (0% response rate; 0 responders out of 22 patients; baseline response rate 37%; odds ratio = 0; Fisher adjusted p-value = .067, Cox-PH adjusted p-value = .33) (23). Moreover, 33 additional markers were significantly associated in only the cox-PH models of progression free survival. Importantly, only 19 of the 56 cohorts exhibited at least one non-responder biomarker association (i.e., Fisher’s test or Cox-PH) (**Supplementary Figure 1c**).

**Figure 1.**
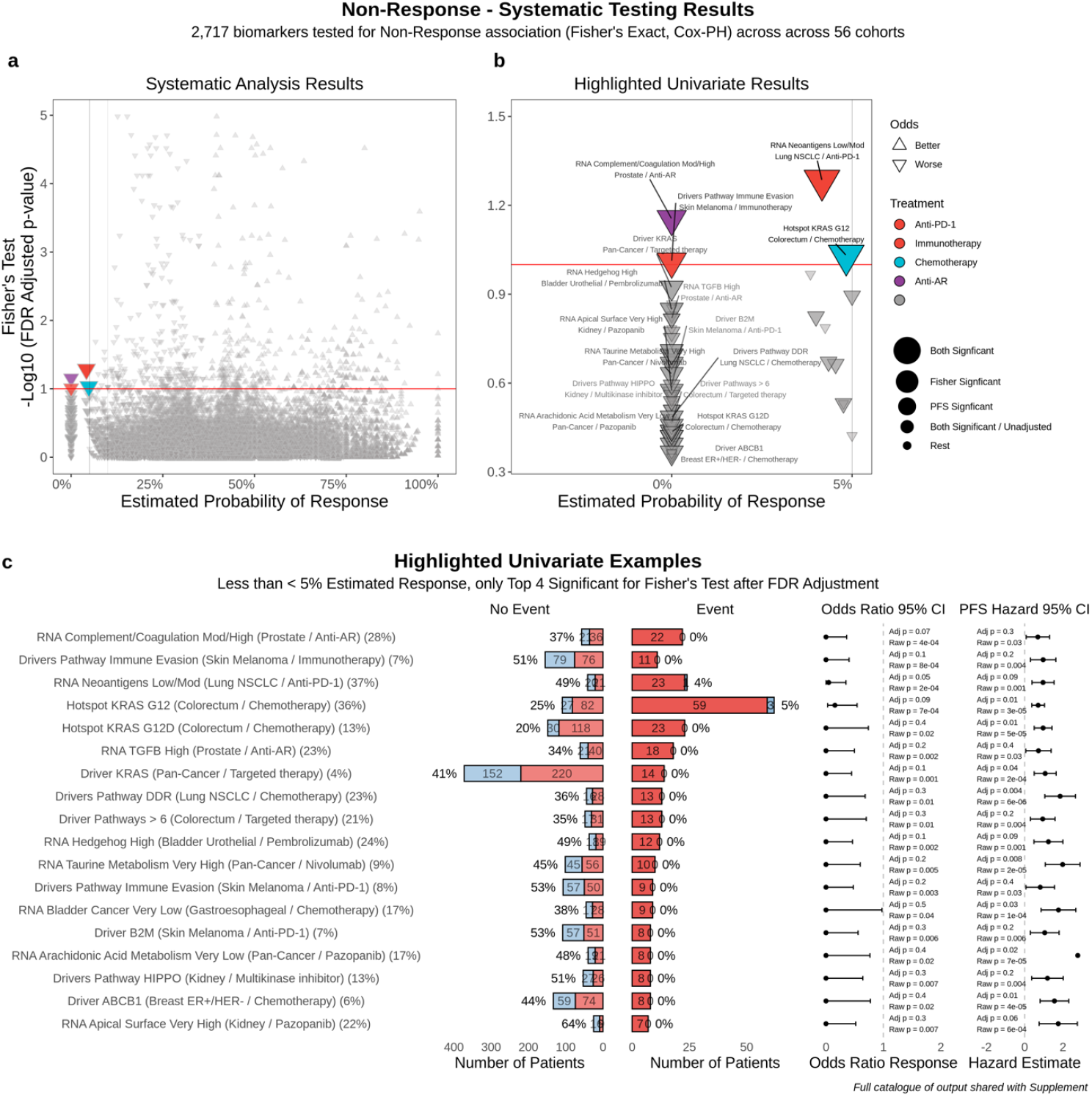
Identification of clinicogenomic biomarkers associated with non-response to therapy across cancer cohorts. **a**, Scatter plot showing the probability of response (x-axis) versus the significance of association with non-response (y-axis) for each clinicogenomic biomarker across treatment and cancer-type cohorts. Markers are color-coded by treatment category. **b**, Zoomed-in view highlighting significantly enriched non-response biomarkers. Vertical and horizontal dashed lines denote thresholds of significance. **c**, Representative examples of non-response biomarkers identified in the analysis. Each row corresponds to one biomarker. Left and right bar plots depict baseline and non-response event response frequencies. Forest plots with 95% confidence intervals indicate odds and hazard ratios.

Notably, many additionally biologically plausible non-responder makers were observed with < 5% response rates, but with statistical significance less than the strict threshold for multiple testing (**Figure 1b-c**). For instance, genomic driver events of the *ABCB1* drug transporter in ER+/HER2− breast carcinoma was associated with a complete lack of response to chemotherapy (0% response rate; 0 responders out of 8 patients; baseline response rate 44%; odds ratio = 0;), consistent with its established role in drug efflux and chemoresistance (24). Similarly, elevated RNA expression of the TGFβ signaling pathway was linked to never-response in patients receiving androgen deprivation therapy (0% response rate; 0 responders out of 18 patients; baseline response rate 34%; odds ratio = 0; Fisher adjusted p-value = .15, Cox-PH adjusted p-value = .36) (25). Importantly, the larger cohorts, especially the pan-cancer ICB cohorts, showed many significant markers enriched for non-response but lacked individual significant markers with estimated chance of response <5% threshold, suggesting a possible lack of true deterministic non-response markers.

The full list of significant associations and their systematic curation are comprehensively annotated in **Supplementary Table 3**, providing a valuable resource that may guide clinical decision and prospective validation in dedicated studies.

### Combined biomarkers to detect non-response to Immune Checkpoint Inhibitors

A challenge from the univariate screening for non-responder markers was that many markers are associated with non-response but still do not define patients with extremely low response rates (< 5%), which are clinically useful. To enable the identification of highly restrictive non-responder biomarkers (i.e., response rate <5% and FDR-adjusted *p* < 0.1), we developed a rational strategy that prioritizes combinations of non-redundant features individually associated with non-response (see **Methods**). As proof of concept, we focused on the immune checkpoint inhibitor treated cohort, which contained the highest number of such biomarkers (see **Methods**). We first selected univariate features significantly associated with non-response based on both Fisher’s exact and Cox proportional hazards models (strict FDR *p* < 0.01). These markers clustered into four biological domains: low TMB, reduced RNA-based T-cell infiltration, elevated RNA expression of renin–angiotensin signaling axis, and high RNA expression of basal cell carcinoma signaling (related to high hedgehog pathway signaling) (**Supplementary Figure 3a-b)**.

We combined these four features into non-response biomarker signature—low TMB, low T cell infiltration, high renin–angiotensin signaling, and high RNA expression of the basal cell carcinoma signature to identify a subset of patients with minimal likelihood of deriving durable clinical benefit and significantly shorter progression free-survival compared to each of the biomarkers individually (0 of 5, 0% response rate, **Supplementary Figure 3c**), suggesting that rational combination of non-redundant biomarkers could be a to further refine subpopulations of patients that will not respond. Furthermore, patients with 3 of these 4 non-response markers also showed a low response likelihood (2/42, 5%) (**Supplementary Figure 3c**). Importantly, this analysis, enabled by a large cohort size and strong univariate associations, highlights the potential of well-powered cohorts to develop robust, multivariate markers of primary resistance.

### Cohort size to detect robust non-response biomarkers

In our systematic study, we highlighted examples of non-responder signals with less than 5% observed response rates across treatment cohorts. However, importantly, the statistical significance of these non-responder signals was assessed by comparison to their respective baseline response rates within treatment cohorts (varying from 18% to 73% in the Hartwig analyses) and not compared directly to the more stringent and relevant low response rate thresholds themselves (e.g. less than 5% or 2%).

Since the ultimate clinical goal of identifying non-response biomarkers is to guide treatment decisions and spare patients from ineffective therapies, we aimed to estimate the highest probability a prospective patient would be a responder when zero responder events are observed for a given biomarker. To this end, we applied the Clopper-Pearson exact method (26) to determine the minimum number of non-responder events necessary to confidently claim patients would have a response rate less than 5% or 2%. Our analysis revealed that at least 59 non-responder events are needed to conclude a response rate below 5% (i.e., upper bound of 95% confidence interval is 5%); while 149 non-responder events are required to establish a response rate less than 2%. The cohort sizes necessary to observe these events depend on biomarker prevalence, with more rare markers requiring larger cohorts. As an example, for the biallelic inactivation of immune escape genes in skin melanoma—where we observed 0% responders, 11 non-response events, 7% biomarker prevalence, 166 patients, and a 51% baseline response rate—the estimated upper bound of the 95% confidence interval is still 24% (**Figure 2a**). Assuming a true 7% biomarker prevalence and 0% observed responders, we estimate that a minimum 890 and 2249 randomly selected melanoma patients undergoing the same treatment would be needed to confidently establish the non-responder’s rate below 5% and 2%, respectively (**Figure 2b**). Taken together, these analyses indicate that current Hartwig datasets, which are among the largest publicly available WGS/WTS datasets with treatment data and clinical responses, are still severely underpowered to detect robust non-response markers that can be used in clinical practice.

**Figure 2.**
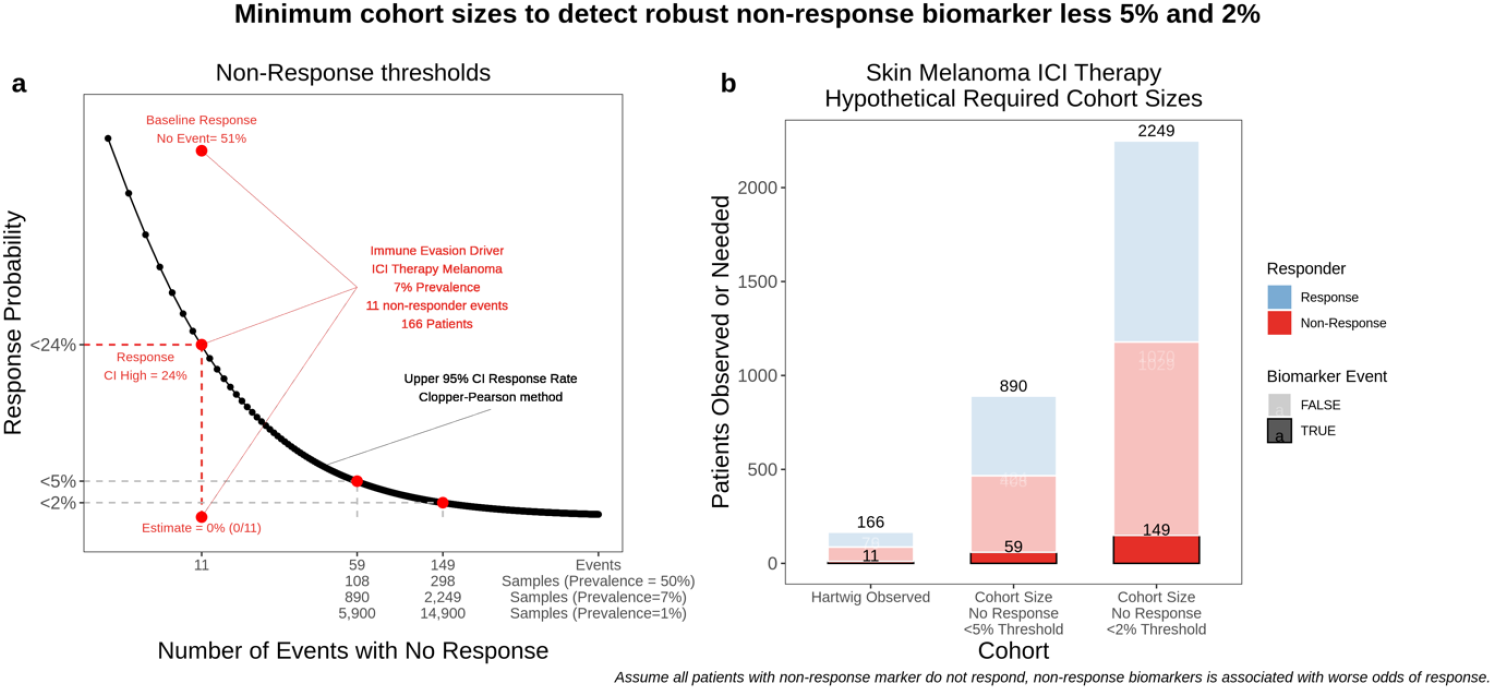
Cohort size estimations for robust non-response biomarkers identification. **a**, Relationship between the number of non-response events (x-axis) and the corresponding response probability (y-axis). Bottom x-axis tick labels show the sample sizes required across different prevalences to reach such a number of non-response biomarkers. Dashed lines mark thresholds for detecting clinically relevant biomarkers occurring at <5% and 2% frequency. Red lines indicate the number of non-response events and the associated response probabilities of immune evasion drivers in the ICI-treated melanoma cohort. **b**, Hypothetical cohort sizes required to detect non-response biomarkers in skin melanoma patients treated with ICIs, shown across different non-response probability thresholds and based on observed biomarker prevalence. ICI, immune checkpoint inhibitors.

## Discussion

Biomarkers of non-response fundamentally differ from stratification biomarkers in that they aim to identify patients unlikely to benefit from specific therapies. The key challenge in evaluating non-response biomarkers is the robust evidence required (e.g., less than 1-5% of probability of response) to justify withholding an otherwise beneficial therapy. This distinction is clinically significant, as reliable non-response biomarkers could help avoid ineffective treatments, reducing unnecessary toxicity and healthcare costs. However, despite their potential, the discovery and clinical implementation of such biomarkers remain challenging.

Robust identification of non-response biomarkers requires large, well-annotated cohorts, as demonstrated by our power analysis. In this study, we conducted a systematic, unbiased screen for biomarkers of non-response using one of the largest integrated resources combining whole-genome and transcriptome sequencing with detailed clinical annotation. While this comprehensive analysis identified several promising candidate biomarkers—many of which are consistent with prior evidence—it also underscored critical challenges for clinical translation. First, the statistical power to confidently define true non-responder subgroups remains limited in a systematic context. In our analysis, only four univariate biomarkers exceeded the stringent significance thresholds. Importantly, statistical power is not solely a function of sample size; factors such as biomarker prevalence and response heterogeneity substantially influence the ability to detect and validate non-response signals. Importantly, the limitation of low biomarker prevalence may be mitigated by aggregating individual driver events into broader molecular pathways—a strategy supported by several of our never-responder signals. Second, many of the identified biomarkers were associated with never-responder phenotypes in relatively small cohorts. Whether these represent true never-responders remains uncertain and requires validation in larger, independent datasets. Third, the scarcity of definitive never-responder biomarkers in larger cohorts—such as those treated with immunotherapy—suggests that combinatorial biomarker strategies may offer improved discriminatory power, as illustrated in our analysis. Finally, it is important to recognize that identifying a biomarker associated with poor response is not equivalent to reliably estimating a sufficiently low probability of benefit—an essential threshold for informing decisions about withholding therapy. Collectively, these findings emphasize the need for expanded, response-annotated real-world datasets and the development of rigorous statistical frameworks to enable the discovery and clinical implementation of non-response biomarkers.

To address these limitations, the incorporation of real-world data from systematic diagnostic profiling efforts could substantially enhance the power and generalizability of non-response biomarker discovery. Equally important is the development of federated data infrastructures that integrate harmonized sequencing protocols and analytical pipelines across institutions and countries. These coordinated frameworks are essential to support large-scale, reproducible identification of non-responder subgroups and to accelerate their clinical translation across diverse healthcare systems.

## Methods

### Hartwig Medical Foundation Database

We used data from the Hartwig Medical Foundation database (DR347, n = 7,011), focusing on a subset of patients with annotated post-biopsy treatment and clinical response data measured using the RECIST criteria (14) (n = 2,596). Clinical data included primary tumour location, treatment information (e.g. drug name, mechanism of action, treatment type), and additional patient metadata such as age, sex, biopsy location, data of biopsy, survival, and pretreatment information.

Whole genome sequencing (WGS) was available for all 2,596 patients, providing rich molecular profiles including somatic mutations, copy number alterations, structural variants (e.g. gene fusions), gene driver events, pathway level gene driver events, homologous recombination deficiency status, HLA status and copy number, telomere length, viral insertions, and V(D)J sequencing output. Bulk RNA-seq data were available for a subset of patients (n = 2,116) and were summarized using computed gene sets and expressed neoepitope predictions.

### Patient Cohorts

Patient cohorts were defined using a combination of primary tumor location, molecular tumor subtype (e.g. Breast ER+/HER-), and post-biopsy treatment characteristics (drug name, mechanism of action, or treatment type). We aimed to balance the trade-off between cohort homogeneity with available sample size (see Supplementary Figure 1b). Cohorts were constructed both across all tumor types (pan-cancer) and within specific cancer types.

A total of 56 cohorts were defined, each containing at least 30 patients and a minimum of 15 responders and 15 non-responders (**Supplementary Figure 1b**). These 56 cohorts included:

- 23 drug specific cohorts (14 pan-cancer, 9 tissue specific)
- 19 drug mechanism cohorts (11 pan-cancer, 8 tissue specific)
- 14 high-level treatment type cohorts (e.g. chemotherapy, targeted therapy, immunotherapy, hormonal therapy; 4 pan-cancer, 10 tissue-specific)

Drug-specific cohorts were designed to find specific resistance signals, while the broader mechanism and treatment-based cohorts leveraged larger sample sizes to identify more generalizable patterns of treatment non-response.

### Derived Biomarkers

We derived a total of 2,594 binary features across clinical, genomic, and transcriptomic domains for systematic association testing. These included 646 gene driver or pathway events, 263 mutational signature features, 88 chromosomal copy number summaries, 68 clinical features, 57 somatic summary metrics, 14 CDR3/VDJ sequences metrics, 12 telomeric length associated features, 8 HLA-related features, 8 markers of genetic immune escape, 8 homologous recombination deficiency features, 1,416 RNA-based gene set expression features, and 6 RNA-based measures of neoepitope burden. All features were binarized based on clinical relevance or quantile-based thresholds. A comprehensive description of feature definitions, thresholds, derivations, and sources is provided in **Supplementary Table 2**.

### Non-Response Definitions and Statistical Analysis

Patient non-response was defined as a binary response using the RECIST clinical response annotations in the Hartwig dataset. Specifically, patients were classified as non-responders if they failed to achieve either a partial response or stable disease lasting longer than six months. To systematically identify features associated with non-response, univariate Fisher’s exact tests were performed with each of the 56 defined patient cohorts. These tests evaluated the association between binary biomarkers and the binary non-response label. Fisher’s exact was selected specifically to handle sparse response patterns, including cases where no response was observed within a feature group. However, a key limitation of the Fisher’s test is its inability to adjust for covariates, and therefore no covariates were included in the testing.

For progression-free survival (PFS) analyses, the hazard event was defined as either RECIST progressive disease or death. Time to event was measured from the date of tumour biopsy to earliest occurrence of a hazard event. For censored patients with no hazard event, the follow-up time was calculated from the biopsy date to the last recorded treatment date or response assessment. For pan-cancer cohorts, primary tumour location was included as a covariate in all models. For RNA-based expression features, tumour purity was included as an additional covariate.

### Multiple Testing Adjustments

To reduce the multiple testing burden and focus on features with higher statistical power, we limited our analysis to those features with at least six total events within a given cohort. In the PFS analyses, features with extreme standard errors due to high collinearity with model covariates were excluded from the results. After these filters, we performed a total of 77,836 successful univariate tests of both Fisher’s exact test and Cox-PH models across the 56 cohorts.

To account for multiple testing, we applied the Benjamini Hochberg (BH) method to control the false discovery rate (FDR), with adjustments performed separately for the Fisher’s exact tests and Cox-proportional hazards models. Given the focus on univariate features with an estimated less than 5% chance of response, large numbers of tests, limited cohort sizes, and sparsity of the biomarker data, and relatively few features passed the multiple testing threshold. Therefore, we chose to highlight features with adjusted p-value < 0.1. While a more conservative FDR correction such as the Benjamini-Yekutieli method would also account for the dependencies among tests, many highlighted features that met the less conservative BH threshold were biologically plausible. Prospective validated analyses would be required for gold standard validation of any highlighted feature.

### Multivariate classifier for Immunotherapy cohort

A key challenge for our systematic non-responder analysis was the stringent focus on univariate features associated with response rates below 5%. Given these strict response rate thresholds, as observed, few features reached statistical significance individually. However, many features showed strong association to non-response across various cohorts, albeit with response rates greater than 5%, suggesting that multivariate combinations of features could be created to better classify non-response.

As an example of a multivariate classifier, we focused on the cohort of patients treated with immune checkpoint inhibitors, of whom 370 had both complete whole genome sequencing and RNA-seq available. From the systematic association testing in this immunotherapy cohort, we selected features the were highly statistically significant for non-response and PFS (Fisher adjusted and Cox proportional hazards adjusted p-values < .01). In total, 17 features passed these significance thresholds, and we performed correlation analyses to identify the independent signals among them.

The correlation analysis showed that these non-responders associated features grouped into four independent clusters with biological meaning:

- Low neoantigen load, low TML, TMB per mega base < 6, low measured RNA neoantigens
- Low T-cell infiltration, 12 gene sets related to reduced immune infiltration.
- High expression of a KEGG gene set for basal cell carcinoma, linked to hedgehog signaling.
- High Expression of a KEGG gene set measuring the renin-angiotensin system.

To build the multivariate classifier, we first selected one representative feature from each cluster (low tumor mutational load, low CD effect T-cell expression, high basal cell carcinoma expression, high renin-angiotensin system expression) and computed for each patient the number of these selected non-response associated features that were present. Patients were then stratified by the number of non-responses associated features, and corresponding response rates along with progression-free survival was compared across these groups (**Supplementary Figure 3)**.

### Minimum Number of Events to Determine Non-Response Thresholds

To derive the minimum number of non-responder events needed to determine whether a patient has less than 5% or 2% of response we used the Clopper-Pearson exact method (26). This method provides a 95% confidence interval for the upper bound of probability of response given no responders were observed. The basic assumption for the Clopper-Pearson exact method is that the number of responses for a given biomarker in a cohort would follow a binomial distribution with the number events equal the cohort size and some fixed unknown underlying probability of response.

## Supplementary Figure Legends

**Supplementary Figure 1.**
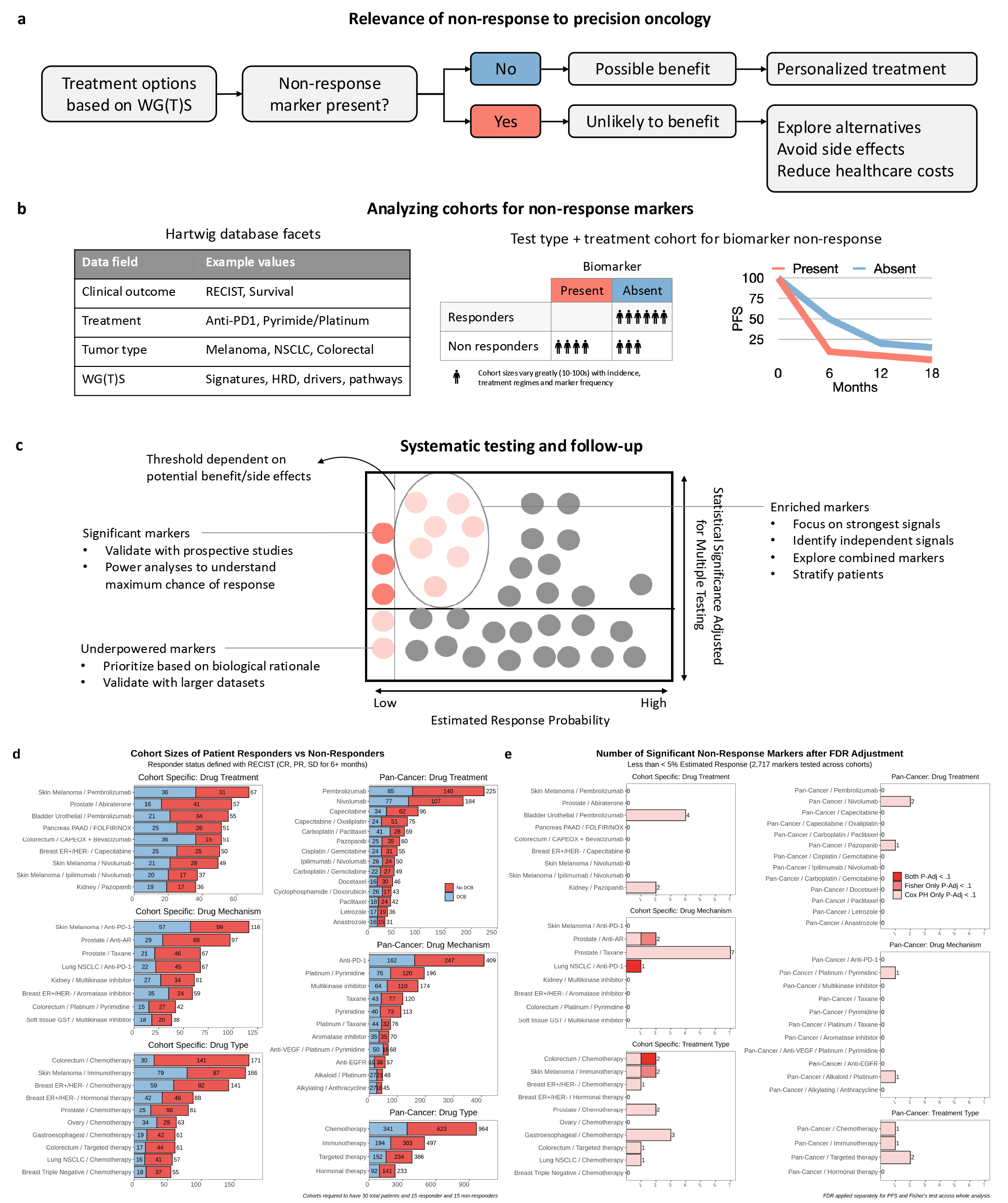
Schematic, Cohort Sizes. **a**, Schematic for motivation to find non-response biomarkers. **b**, Use the Hartwig database to derive clinical outcomes, define cohorts, and create binary biomarkers. Each biomarker is tested systematically for each cohort with Fisher’s exact test for association to non-response and with cox-proportional hazards models for progression free survival. **c**, propose ways systematic results can be prioritized for 1) significant non-response markers, 2) under-powered non-response markers, and 3) markers enriched for non-response. **d**, Names and number of patients for the 56 defined cohorts. **e**, Total FDR-adjusted significant non-response markers for fisher’s exacts tests and cox-proportional hazards models.

**Supplementary Figure 2.**
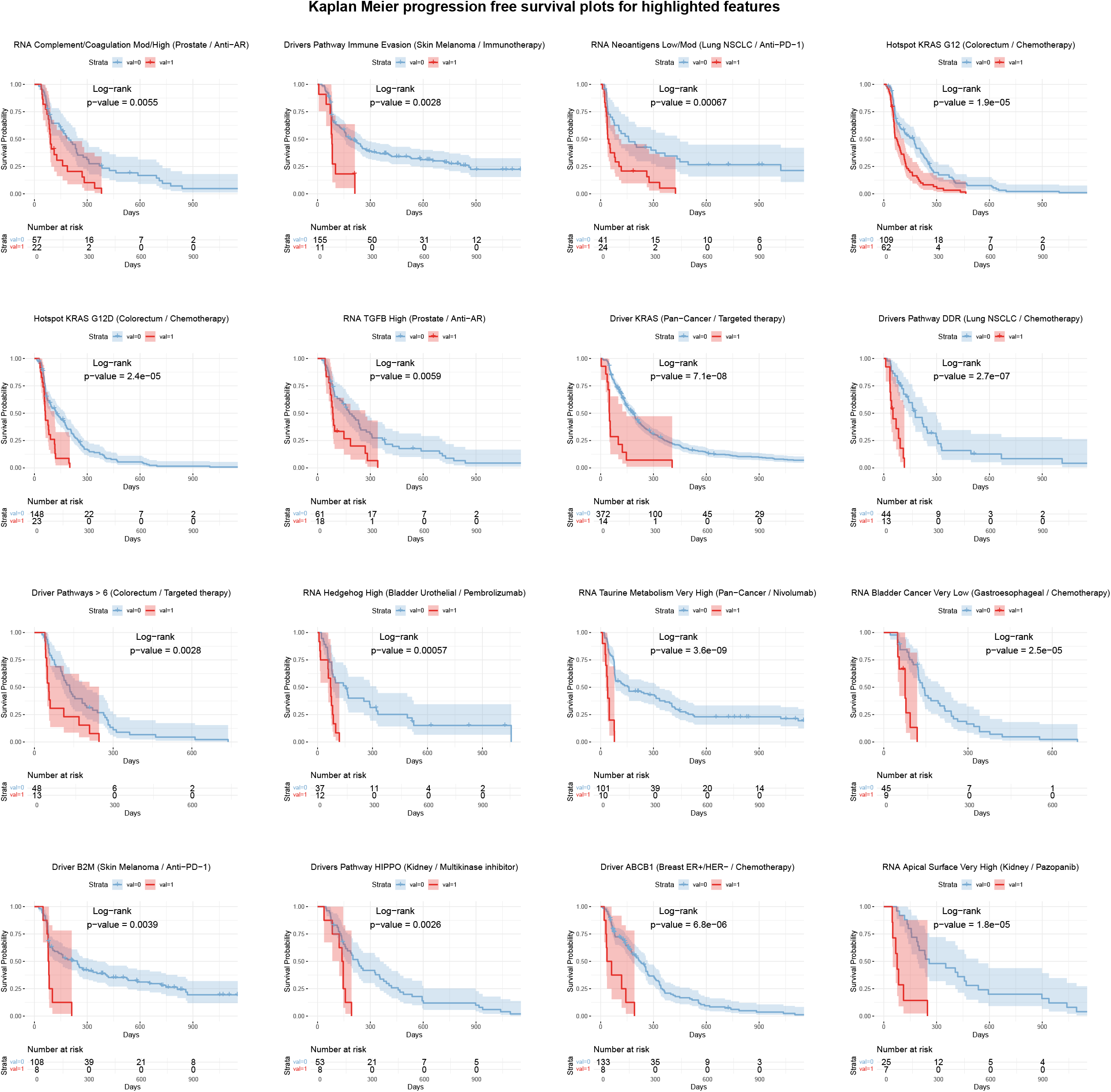
Kaplan Meier progression free survival plots for highlight features. **a**. Kaplan-Meier curves of progression-free survival (PFS) for patients stratified by the presence (red) or absence (blue) of the respective non-response associated and highlighted feature. Reported p-values correspond to the log-rank test comparing the PFS between the presence or absence biomarker groups.

**Supplementary Figure 3.**
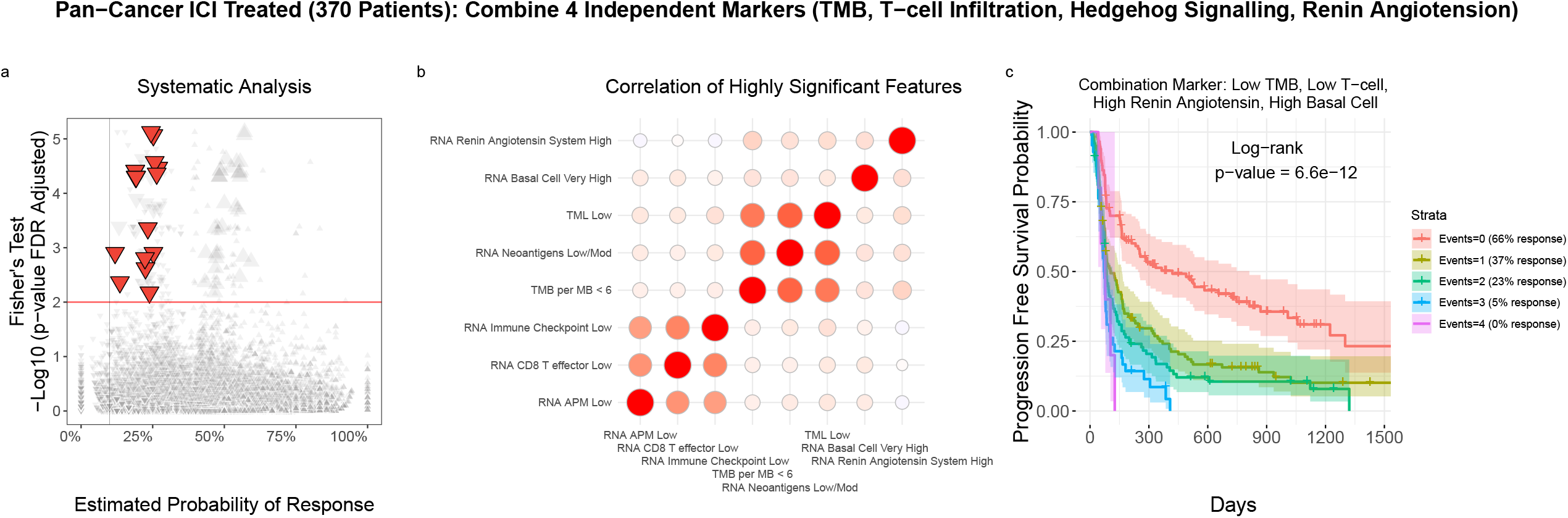
Combination Marker for immune checkpoint inhibitor treated cohort. **a**, Scatter plot showing the probability of response (x-axis) versus the significance of association with non-response (y-axis) for each clinicogenomic biomarker across treatment and cancer-type cohorts. FDR adjusted significant markers (p-value < .01) associated with both non-response and poor PFS survival outcomes in the immunotherapy cohort are highlighted and enlarged in red. The down arrows indicate each marker is associated with worse odds of response. **b**, Correlation matrix of selected non-responder markers from the systematic analysis, showing four clusters of independent features. **c**, Kaplan Meier plots showing progression free survival in the 370 immunotherapy-treated patients, grouped by the number of non-responders associated biomarkers observed for each patient (0 to 4).

## Data availability

Access to the genomic, transcriptomic, and clinical metadata from the Hartwig Medical Foundation data can be obtained freely through data access requests to the Hartwig database (https://www.hartwigmedicalfoundation.nl/en/data/data-access-request).

## Code availability

The source code to reproduce the analysis for the manuscript is available in the following repository: https://github.com/Computational-Immunogenomics/hmf_biomarkers

## Acknowledgements

This publication and the underlying study have been made possible partly on the basis of the data that Hartwig Medical Foundation and the Center of Personalized Cancer Treatment (CPCT) have made available to the study. F.M.J. acknowledges support from a Ramón y Cajal Fellowship (RYC2022-037005-I) funded by the Spanish Ministry of Science. This work was also supported by the FERO Foundation (XXIV Beca FERO) and by the CaixaResearch Advanced Oncology Research Program supported by “La Caixa” Foundation. VHIO would like to acknowledge the Cellex Foundation for providing research facilities and equipment and the State Agency for Research (Agencia Estatal de Investigación) the financial support as a Center of Excellence Severo Ochoa (CEX2020-001024-S/AEI/10.13039/501100011033).

## Author contributions

E.C. and F.M.-J. conceptualized the study. J.U., J.L., S.R., E.C., and F.M.-J. developed the methodology. J.U., J.L., and S.R. provided software. J.U., J.L., S.R., P.R., E.C., and F.M.-J. validated the results. J.U., J.L., and S.R. conducted the formal analysis. J.U., J.L., S.R., and F.M.-J. performed the investigations and provided resources. J.U., J.L., S.R., P.R., F.M.-J. curated the data. J.U. and F.M.-J. wrote the original draft of the manuscript. J.U., J.L., S.R., P.R., E.C., and F.M.-J. reviewed and edited the manuscript. J.U., J.L and F.M.-J. performed visualization. J.L., E.C., and F.M.-J. supervised the project. J.L., E.C., and F.M.-J. managed project administration. E.C. and F.M.-J. acquired funding.

