## Supplementary Note 1 for "Systematic identification of genomic non-response biomarkers to cancer therapies"

### B2M Biallelic Loss Validation

Broad validation of the systematic study is challenging due to diverse external datasets and studies, which differ in patient populations, molecular data formats and availability, and levels of clinical metadata and response annotation.

As an example, we performed a validation analysis focusing on the biallelic B2M loss in metastatic melanoma patients treated with checkpoint inhibitor therapies. To this end, we used publicly available studies accessed through cBioPortal (1).

From cBioPortal, we selected the three available immunogenomic studies of checkpoint inhibitor therapies in metastatic melanoma (2–4). Each study captured B2M biallelic deletions and included annotated clinical patients' responses measured using the RECIST criteria and overall survival. The treatment mechanism for two larger studies was Anti-CTLA4, while one study contains Anti-PD1 treated patients. This mix of treatment mechanisms differs somewhat from the Hartwig cohort which contains mostly Anti-PD1 or Anti-PDL1 treated patients.

Across the three studies, 210 total metastatic melanoma patients were available, with 6 patients showing B2M biallelic loss, indicating a slightly lower prevalence compared to the Hartwig cohort. Patient breakdown by study is as follows:

- Snyder et. al (Anti-CTLA4, 64 total patients, 3 patients with biallelic B2M Loss, all labeled non-responsive to treatment, but one labeled LB in durable clinical benefit)
- Van Allen et. al (Anti-CTLA4, 110 total patients, 3 patients with biallelic B2M loss, all labeled Progressive Disease for RECIST)
- Hugo el. Al. (Anti-PD1, 38 patients, no measured biallelic B2M loss)

In total, all six patients with biallelic B2M biallelic loss across these studies were labeled as either non-responsive or having progressive disease. One patient in the Snyder et al. study was labeled to be non-responsive but also have LB for durable clinical benefit, however, this patient showed the worst overall survival outcome of any patient with this LB label. These studies underscore the challenge of validation of rare biomarkers such as B2M loss that are observed only rarely in study cohorts.

We also performed an overall survival analysis using data from the cBioPortal studies. First, we compared the 6 patients with biallelic B2M loss to 204 patients without B2M biallelic loss. In this marginal analysis, the Cox proportional hazards model estimated an increased log-hazard of .48 for patients with B2M biallelic loss (non-significant p = 0.29; **Supplementary Note Figure 1a).** To further investigate, we stratified the analysis by treatment response. When we compared the biallelic B2M loss patients to the other 128 non-responders, the estimate log-hazard was similar (.08, p=0.86), indicating no observed meaningful difference. However, in comparison to the 68 responders, patients with biallelic B2M loss showed a much higher estimated log-hazard of 1.27, which was statistically (p = 0.009; **Supplementary Note Figure 1b).**

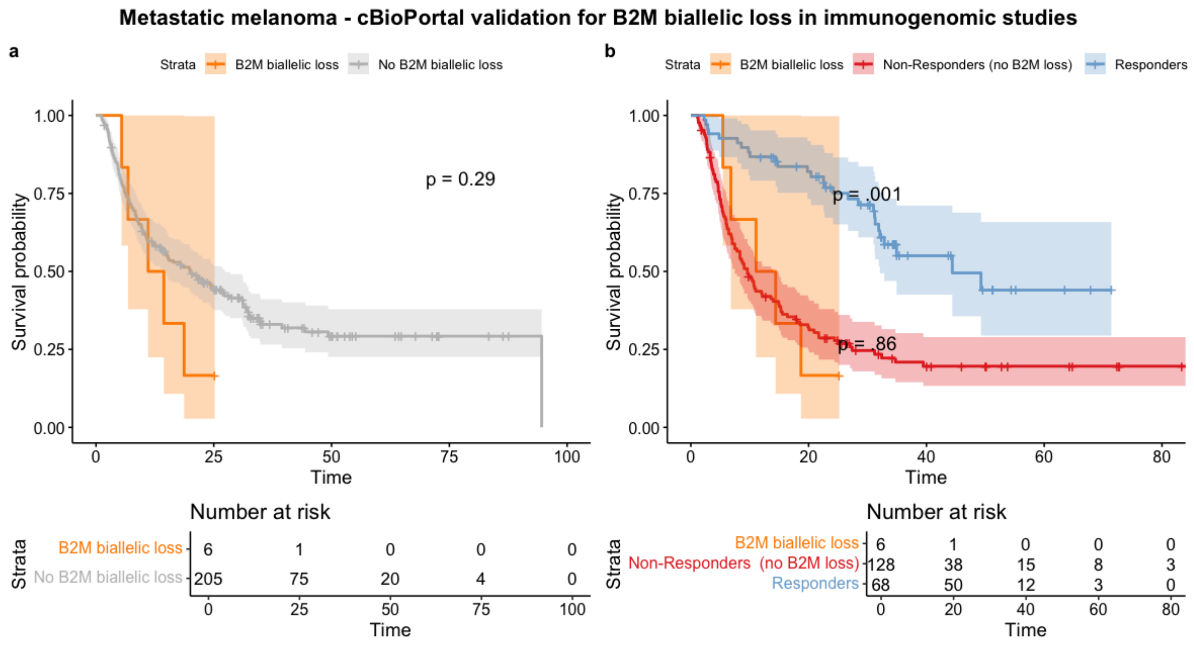


**Supplementary Note Figure 1. a,** Kaplan Meier plots comparing overall survival for the 6 patients with B2M biallelic loss versus 205 patients without B2M biallelic loss across 3 selected metastatic melanoma studies with B2M loss information from cBioPortal. **b,** Kaplan Meier plots comparing overall survival for patients with B2M biallelic loss versus patients without B2M biallelic loss further stratified by whether they were labeled as either responder or non-responder in their respective clinical study.
